# Invasive European green crab (*Carcinus maenas*) predation revealed with quantitative DNA metabarcoding

**DOI:** 10.1101/2023.11.27.568928

**Authors:** Mary C Fisher, Emily W. Grason, Alex Stote, Ryan P. Kelly, Kate Litle, P. Sean McDonald

## Abstract

Predation by invasive species can threaten local ecosystems and economies. The European green crab (*Carcinus maenas*), one of the most widespread marine invasive species, is an effective predator associated with clam and crab population declines outside of its native range. In the U.S. Pacific Northwest, green crab have recently experienced increases in abundance and expanding distributions, generating concern for estuarine ecosystems and associated aquaculture production. However, regionally-specific information on the trophic impacts of invasive green crab is highly limited. We compared the stomach contents of green crabs collected on aquaculture beds versus natural intertidal sloughs in Willapa Bay, Washington, to provide the first in-depth description of European green crab diet at a particularly crucial time for regional management. We first identified putative prey items using DNA metabarcoding. For key prey species, we also applied a quantitative model to calculate the true abundance of each species in stomach content DNA samples. From the stomach contents of 62 green crabs, we identified 56 unique taxa belonging to nine phyla. The stomach contents of crabs collected from aquaculture beds were significantly different from the stomach contents of crabs collected at natural intertidal sloughs. Across all sites, arthropods were the most frequently detected prey, with the native hairy shore crab (*Hemigrapsus oregonensis*) the single most common prey item. Of the eight species included in the quantitative model, two ecologically-important native species – the sand shrimp (*Crangon franciscorum)* and the Pacific staghorn sculpin (*Leptocottus armatus*) – were the most abundant in crab stomach contents, when present. In addition to providing timely information on green crab diet, our research demonstrates the novel application of a recently developed model for quantitative DNA metabarcoding. This represents another step in the ongoing evolution of DNA-based diet analysis towards producing the quantitative data necessary for modeling invasive species impacts.

## Introduction

Impacts of biological invasions can be difficult to measure at the early stages for a variety of reasons. In many cases, managers lack baseline data against which to compare population changes in the local ecological community (Lindenmayer & Likens, 2010). Even when such data do exist, per capita impacts may not scale up to statistically detectable population community shifts when the overall invasive population size remains relatively small (Parker et al., 1999). Moreover, similar to detection and establishment, assessment of impacts can “lag” behind their occurrence if impacted species have not yet had enough time to incur detectable demographic shifts, e.g., changes in vital rates (Crooks, 2002; Kolar & Lodge, 2001). Yet gauging impacts early in the invasion process can buy managers valuable time in prioritizing limited resources to geographies or habitats before they incur dramatic losses once the invasive species has reached greater abundances (Mehta et al., 2007).

For predatory invasive species, diet studies offer crucial early insight into the range and magnitude of trophically-mediated community impacts. Diet analysis can also be combined with manipulative experiments and behavioral studies to parameterize population-level impact analysis for later invasion stages. For example, several diet studies of the invasive lionfish (*Pterois volitans* and *P. miles*) elucidated the range, electivity, and size-dependency of lionfish diets in the Caribbean (Eddy et al. 2016; Morris and Akins 2009; Layman and Allgeier 2012; and others cited in McCard et al. 2021); when results were combined with behavioral observations and models on diurnal activity patterns (Green et al., 2011), prey switching (McCard et al., 2021) and functional responses (DeRoy et al., 2020), researchers were able to predict population-level impacts.

While diet analysis has traditionally relied on visual identification of stomach contents to directly characterize predation (including Eddy et al. 2016, Morris and Akins 2009, Layman and Allgeier 2012), DNA metabarcoding is an increasingly popular method to describe a predator’s diet. DNA metabarcoding has been shown to identify stomach contents to higher taxonomic resolutions than visual analysis, and can reveal novel predator-prey interactions through improved detection of certain prey groups, that are visually indistinguishable after mechanical digestion (e.g., those composed primarily of soft tissue; Berry et al., 2015; Dahl et al., 2017; Nielsen et al., 2018). There have also been recent methodological improvements in the analysis of DNA metabarcoding data more broadly, aimed at the difficulty of translating metabarcoding data – which consists of DNA sequencing read abundances – into accurate quantitative information about the sampled environment. Chief among these challenges is accounting for bias introduced during the required DNA amplification process, which causes sequencing read abundance to be unrepresentative of the true composition of a DNA sample (Kelly et al., 2019; Silverman et al., 2021). However, quantitative modeling methods that were recently developed for other applications of DNA metabarcoding can now be paired with the construction of mock prey “communities” to calibrate sequencing read abundance and estimate the true composition of DNA extracted from stomach contents.

We combine DNA metabarcoding of crab stomach contents with quantitative models to describe the diet of invasive European green crabs (*Carcinus maenas*; hereafter “green crab”) in Willapa Bay, Washington, where there have been recent dramatic increases in crab abundance. Green crabs were first detected in Willapa Bay in 1998 (Dumbauld & Kauffman, 1998), but apparently failed to establish a sustained presence in the estuary during the subsequent two decades (Behrens Yamada et al., 2022). Recent trapping and monitoring efforts, as well as opportunistic observations made by local shellfish growers, detected a dramatic increase in both the relative abundance of green crabs as well as their geographic range within the Bay ca. 2015 – 2017. These trends have raised concern that shifting conditions are enabling green crabs to transition from a lag phase into population growth, establishment, and spread. These later phases of the invasion process are most commonly associated with impacts to native and commercially important resident species. Regional resource managers anticipate significant impacts of the green crab invasion, with particular concern for the native habitat-building eelgrass (*Zostera marina*), and shellfish species of high commercial and sociocultural importance. Willapa Bay contributes approximately a quarter of all Washington state shellfish aquaculture production (Washington Sea Grant, 2015), predominantly through the harvest of Manila clam (*Ruditapes philippinarum*) and Pacific oyster (*Magallana gigas*; Washington Sea Grant, 2015). Shellfish growers have expressed concern that green crab might already be reducing the recruitment of natural set Manila clams on which the industry relies.

Though species- and community-level impacts of green crab have been documented in other parts of their invasive range (Glude, 1955; Grosholz et al., 2000; de Rivera et al., 2011), most remain speculative in Washington based on likely or observed habitat overlap between green crab and species that have been impacted elsewhere, or on behavioral experiments that might not translate to population impacts in the field. For instance, McDonald et al.(2001) demonstrated that juvenile green crab can both prey on and outcompete size-matched native Dungeness crab (*Cancer magister*) for both food and shelter, but the two species did not co occur at that size during that period. Additionally, while previous research has documented declines in the native hairy shore crab (*Hemigrapsus oregonensis*) attributable to green crab in California (Grosholz et al., 2000; de Rivera et al., 2011), and the two species overlap extensively in Washington (WSG Crab Team), no baseline data exist to robustly evaluate local green crab impacts to *H. oregonensis* populations. Yet, feeding behavior of generalist invasive predators can vary substantially across their global range (Glassic et al., 2023; McAulay et al., 2021). Thus, local, habitat-specific diet analyses are needed to assess the variability and context dependence of diets and consequent community impacts.

We provide a timely investigation of European green crab diet at aquaculture sites and natural sloughs in Willapa Bay, Washington, and quantify the consumption of prey species of interest within and across site types. We first identified putative prey taxa from green crab stomach contents using DNA metabarcoding of the mitochondrial cytochrome oxidase I (COI) region. We then constructed mock prey “communities” for a subset of species of particular socioeconomic and ecological concern, and used a novel quantitative model (Shelton et al., 2023) to correct for amplification bias and estimate the true abundance of each species’ DNA in crab stomach contents. We used this abundance data to model the composition of the average diet of a green crab from an aquaculture site versus natural slough. Our work not only provides the first detailed description of *in situ* green crab diet in Washington State, it also broadens the potential downstream applications of diet DNA (dDNA) to invasive species research and management by incorporating novel quantitative methods from the broader environmental DNA literature.

## Methods

### Sample collection

We collected green crabs at four locations in Willapa Bay, Washington, in May, July, and September of 2021 (Fig 1, Table 1). Two collection sites were located on tide flats under active cultivation of un-netted Manila clams (*Ruditapes philippinarum*; “clam bed” or aquaculture sites), and two were natural intertidal sloughs (“slough” sites), adjacent to emergent salt marsh vegetation. Sites were paired such that one of each type was found on either side of a transition zone within Willapa Bay known as the “fattening line.” Sites to the north of the fattening line (toward the estuary mouth), are characterized by greater influence of ocean water, with lower residence time than those to the south (toward the head of the estuary; Banas et al. 2007). Spring and summer months were sampled under the assumption that temperature-influenced energetic demands increase foraging activity (Aagaard et al., 1995; Elner, 1980), potentially reducing the incidence of empty stomachs and increasing encounter rates with prey of interest.

**Figure 1.**
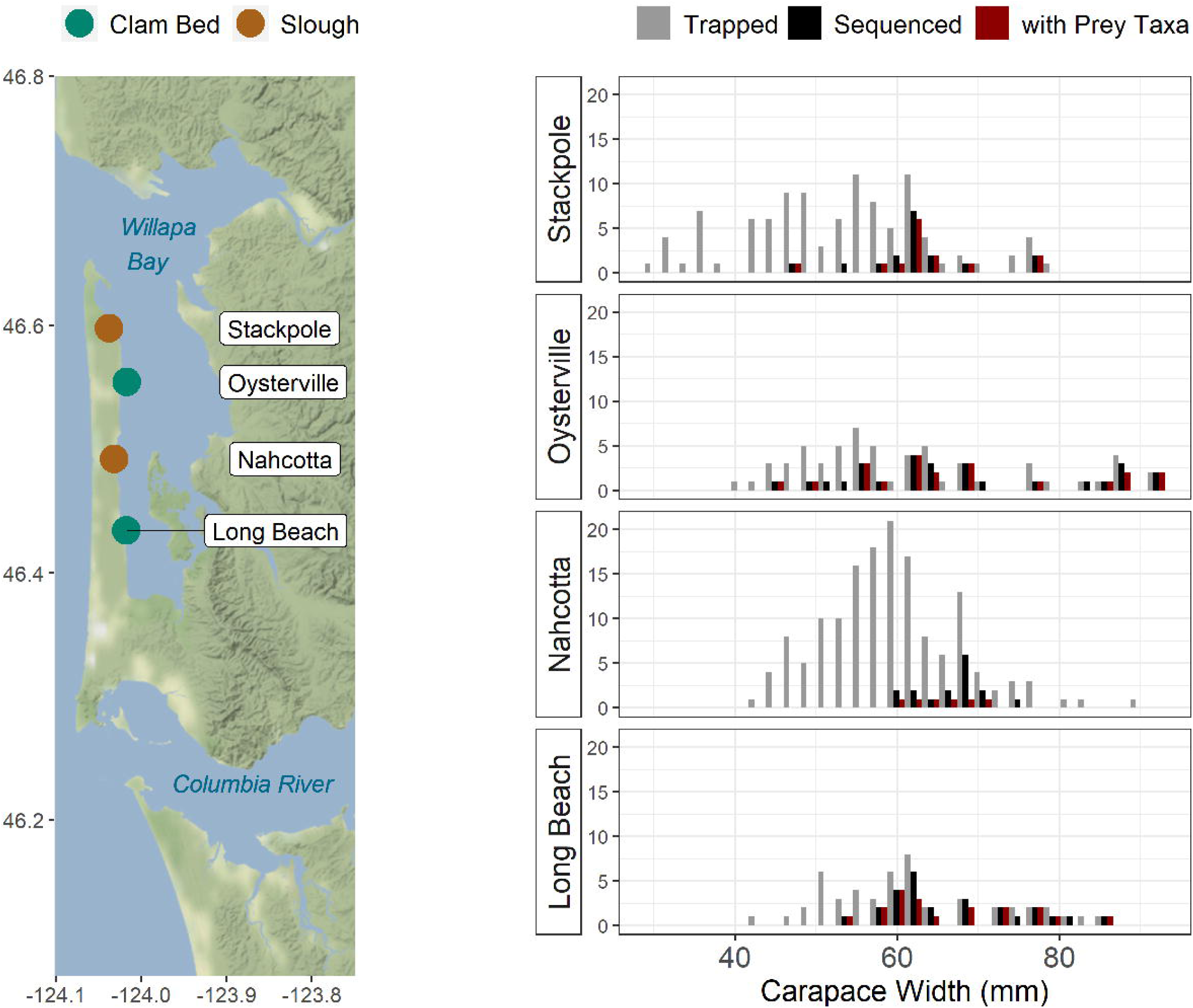
Map of the four collection sites in Willapa Bay, WA, with the distribution of carapace widths (in millimeters) at each site for all European green crab trapped (gray), those chosen for sequencing (black), and then those chosen for sequencing which had identifiable prey DNA in their stomach contents.

**Table 1.** Collection sites, dates, and sample sizes, with collection-specific information on crab and prey. The final sample size (“Final N”) is the number of sequenced European green crab with putative prey items identified from stomach contents. We also provide the median crab carapace width (mm) and the overall prey alpha diversity for the given site.

| Site Type | Site Name | Site Latitude, Longitude | Sampling Dates | N Dissected | N Sequenced | Final N (% Sequenced) | Mean Carapace Width (mm) | Prey $\alpha$ Diversity |
| --- | --- | --- | --- | --- | --- | --- | --- | --- |
| Slough | Stackpole | 46.597556, -124.037698 | 8 Jul 2021<br>7 Sep 2021 | 25 | 17 | 14 (0.82) | 62.5 | 16 |
| Slough | Nahcotta | 46.492106, -124.031611 | 8 Jul 2021<br>7 Sep 2021 | 26 | 16 | 6 (0.38) | 66.9 | 9 |
| Clam Bed | Long Beach | 46.434368, -124.017641 | 26 May 2021<br>8 Jul 2021<br>7 Sep 2021 | 34 | 26 | 20 (0.77) | 66.8 | 20 |
| Clam Bed | Oysterville | 46.554476, -124.01738 | 26 May 2021<br>8 Jul 2021<br>7 Sep 2021 | 36 | 27 | 22 (0.81) | 67.8 | 39 |

Crabs were captured using galvanized steel minnow traps and square Fukui fish traps, baited with Pacific mackerel (*Scomber japonicus*) and set for an overnight high tide (for detailed trapping protocol, see Supplementary Material). On trap retrieval, green crabs were sexed and their carapace width measured (to the nearest millimeter), and immediately placed on dry ice to preserve stomach contents from degradation. Crabs were transported to the University of Washington on dry ice, and stored in a -20C freezer until dissection. Of the 344 total green crabs captured during these efforts, 121 were selected for dissection. We preferentially selected larger, predominantly male crabs for analysis to maximize the likelihood that crabs would be large enough to handle mature adult Manila clams (Fig. 1).

We removed stomach contents from each crab by inserting a pipette with a filter tip through each crab’s mouth and esophagus, in order to limit the amount of predator DNA and gut bacteria included in the stomach sample for DNA extraction (following Cordone et al. 2022). If the stomach was nearly empty, we used gastric lavage (with 400uL 100% EtOH) to help remove particulates. Stomach contents were deposited in a sterilized 1.5mL microcentrifuge tube and stored in 100% EtOH at -20C until DNA extraction. Stomach fullness was then ranked on a 0 (empty) to 7 (full) scale (Stevens et al., 1982). All dissection tools were sterilized between individuals with 20% NaClO, 100% EtOH, and flame.

### DNA extraction and metabarcoding

DNA was extracted from stomach contents with the Qiagen DNEasy Blood & Tissue Kit (Qiagen Corp., Valencia, CA, USA), with some modifications to the kit protocol, including an initial drying period in a vacuum centrifuge to remove 100% EtOH. A subset of samples did not yield adequate DNA from the DNEasy Blood & Tissue Kit, and so were re-extracted using the Qiagen PowerSoil Pro Kit (Qiagen Corp., Valencia, CA, USA).

DNA metabarcoding was completed using a two-step PCR protocol (modified from Jacobs-Palmer et al., 2021). The first PCR step used the BF3/BR2 primer pair to amplify a 418bp segment (within the Folmer fragment) of the mitochondrial Cytochrome Oxidase I (COI) region (Cordone et al., 2022). We chose this primer pair because of its prior application in green crab diet analysis (Cordone et al., 2022), and because preliminary analysis on a subset of our samples showed its ability to amplify DNA from prey species of particular management and conservation interest. The 25uL PCR reaction consisted of 2X Qiagen Multiplex Plus Master Mix, 0.5uM of each primer, and 4uL of DNA extract. After an initial denaturation step (95°C for 5 minutes), we ran the annealing sequence for 35 cycles (95°C for 30 seconds;, 50°C for 30 seconds; 72°C for 50 seconds), followed by a final extension at 68°C for 10 minutes. All DNA extracts were amplified in three separate reactions to produce three technical replicates per crab stomach. PCR product was visualized on a 1.5% agarose gel, cleaned to remove primer dimers using Mag-Bind Total Pure NGS beads (Omega Bio-Tek, Norcross, GA, USA), and quantified using Quant-iT dsDNA High Sensitivity Assay Kit (ThermoFisher Scientific, Waltham, MA, USA). We then diluted all samples with high enough concentrations of cleaned amplicon to 10ng DNA in 11.25uL by adding the appropriate amount of UltraPure DNase/RNAse-free water (Fisher Scientific, Pittsburgh, PA, USA). The second PCR added index sequences (6 base pair nucleotide tags; IDT for Illumina DNA/RNA UD Indexes) to the amplicons using 1X HiFi HotStart Ready Mix (KAPA Biosystems, Wilmington, MA, USA) and 11.25uL of diluted (high concentration samples) or pure (low concentration samples) amplicon. The number of PCR cycles varied between 5, 8, and 12 depending on amplicon concentration. Indexed amplicons were visualized on a 1.5% agarose gel, cleaned, and quantified using the same procedures as the first PCR step. This process yielded at least three laboratory samples per crab of uniquely indexed amplicons.

The same two-step PCR process was completed for a PCR positive control (a DNA sample from kangaroo, genus *Macropus*), a PCR negative control, and a DNA extraction negative control. Positive and negative controls from across DNA extractions / PCR runs were pooled prior to indexing, so that each sequencing run contained one positive control sample and one negative control sample.

We pooled indexed amplicons and controls for 2×300 (paired end) sequencing on an Illumina MiSeq v3. The first sequencing run included indexed amplicons from n=30 crab (90 total samples) plus controls, and was completed by the University of California, Los Angeles’ Technology Center for Genomics & Bioinformatics. The remaining three sequencing runs consisted of indexed amplicons from n=56 samples (168 technical replicates) plus controls, and was prepared and sequenced in-house using the MiSeq Reagent Kit v3 600-cycle (Illumina, San Diego, CA, USA) at the Center for Environmental Genomics Laboratory, University of Washington. All pooled libraries sequenced at the University of Washington’s Center for Environmental Genomics were run at a concentration of 7pM with 20% phiX.

### Prey identification

DNA sequencing data was cleaned and analyzed in R v4.1.3 and git bash v4.4.19, using a combination of custom scripts and the programs cutadapt (Martin, 2011), DADA2 (Callahan et al., 2016), and NCBI’s BLAST, to translate demultiplexed DNA sequencing data into a filtered dataset of putative prey Amplicon Sequence Variants (ASVs) with corresponding sequence counts (modified from Gallego et al., 2020). In short, we trimmed primers from demultiplexed DNA sequences using cutadapt run with git bash, and trimmed sequence length according to base pair read quality with DADA2 in R. We then used DADA2 to filter the trimmed sequencing data with the default parameters (maximum of two expected errors, reads truncated at first occurrence of a quality score of two); and to identify ASVs in the filtered sequences, with a parametric error model built on the sequencing data itself (Callahan et al., 2016). We also used DADA2 to identify and remove chimeric ASVs, and then removed ASVs found in the positive or negative controls. We aligned the filtered ASVs to NCBI’s BLAST database using blastn, with a minimum percent identity of 94%, minimum word size of 15, and maximum e-value of 1e-30. We identified the last common ancestor for any ASVs that returned multiple species from blastn.

Additional filtering was completed with custom scripts to remove ASVs that matched to green crab in blast, as we are unable to distinguish between predator DNA and putative prey of the same species; and taxa considered non-target (e.g., bacterial, fungal taxa) or too small to be targeted prey (e.g., diatoms, rotifers, unicellular algae, etc.). We first retained only ASVs that were identified to species, and then added higher taxonomic identifications back into the data if they were not already represented in the species dataset for a given crab.

From this processed taxonomic dataset, we used a process modified from Duprey et al. (2023) to manually categorize each taxon as a (1) native taxon, (2) previously detected non-native taxon, or (3) previously undetected non-native taxon. If the Washington coast was included within the taxon’s indigenous distribution in one or more of the World Register of Marine Species (WORMS), the Biodiversity of the Central Coast database, the National Estuarine and Marine Exotic Species Information System (NEMESIS), the Encyclopedia of the Puget Sound, or a peer-reviewed primary source, the taxon was considered native. Taxa were considered “previously detected non-natives” if there was a known invasion history described in any database. A taxon was considered a “previously undetected non-native” if none of the databases included the Washington coast in the taxon’s known distribution. Any taxon labeled non-native was then subject to additional filtering, to account for the possibility that ASVs for uncatalogued or poorly represented native taxa to be assigned to non-native close relatives. For previously detected non-natives: if the percent identity with the BLAST reference sequence was >95%, or if percent identity was <95% but native sister taxa were represented in the BLAST database, the taxon was retained; otherwise, we either retained the genus shared by the matched non-native taxon and any native sister taxa (for species-level identifications), or we removed the taxon from the final dataset. For previously undetected non-natives: if the percent identity with the BLAST reference sequence was >98% and native sister taxa were represented in the database, the taxon was retained (none met this criteria); otherwise, as above, we either retained the identification to the genus level or removed the taxon from the final dataset.

The final filtered dataset consisted of 56 unique taxa associated with 266 total ASVs. Although the DNA from these taxa could be present in the stomach contents for a number of reasons not limited to targeted predation, which we address in the discussion, we will hereafter refer to these taxa as “prey” that comprise the green crab diet.

### Analyzing diet variability

We used presence/absence information from the final prey dataset to calculate the α diversity of the stomach contents per site, and β diversities between sampling sites (using the R package vegan; Oksanen et al., 2022). To determine whether diet composition differed significantly among the four sampling sites, we conducted a PERMANOVA and then used a PERMDISP, from the *adonis2* function in vegan, to evaluate whether the significance in the PERMANOVA was due to variability in sample dispersion around centroids as opposed to differences in centroid location (Oksanen et al., 2022). We also identified which site pairs had significantly different prey assemblages by conducting post-hoc pairwise comparisons with permutation tests implemented in the R package *pairwiseAdonis* (Martinez Arbizu, 2017). We completed all statistical testing first using a presence/absence matrix and the Jaccard dissimilarity index, and then using the eDNA index, an index of read-count proportions (Kelly et al., 2019), and the Bray-Curtis distance (Guri et al., 2023). All code is archived on Github (mfisher5/Green-Crab-dDNA-Willapa-Bay).

### Quantifying key prey contributions to stomach contents

While the eDNA index allows us to compare sequence read abundances across sites within an individual taxon, bias introduced during PCR processes means that read abundances cannot be directly compared across different taxa (Kelly et al., 2019; Shelton et al., 2023). This limits our interpretation of the relative impact of green crab on different prey items of particular ecological or socioeconomic importance. To overcome these limitations for a subset of prey of interest, we converted proportions of sequenced amplicons (read abundances) to proportions of DNA template (true abundance) using a calibration sequencing run of “mock communities” and a quantitative model (Shelton et al. 2023). This model corrects read proportions in DNA metabarcoding data for primer- and species-specific amplification efficiency. Posterior mean estimates from the model represent the proportion of DNA from each prey taxon that was present in the PCR template (i.e., subsampled DNA extract from green crab stomach contents).

The Shelton et al. (2023) model requires sequencing data from “mock community” samples that consist of known concentrations of DNA from each species of interest. To construct mock communities, we collected tissue samples from nine prey species, chosen for presence as bycatch in green crab traps, management or conservation interest, and their frequency in the stomach content DNA metabarcoding data (hereafter, “sample data”): Manila clam (*Ruditapes philippinarum*), soft-shell clam (*Mya arenaria*), Dungeness crab (*Cancer magister*), hairy shore crab (*Hemigrapsus oregonensis*), sand shrimp (*Crangon franciscorum*), invasive mud snail (*Batillaria attramentaria*), eelgrass (*Zostera marina*), shiner perch (*Cymatogaster aggregata*), Pacific staghorn sculpin (*Leptocottus armatus*), and prickly sculpin (*Cottus asper*). We also included predator (green crab) DNA. DNA was extracted from tissue samples using the Omega Bio-Tek EZ-DNA Mollusc Extraction Kit (Omega Bio-Tek, Norcross, GA, USA). DNA extracts were amplified using the same PCR process as for the stomach content samples, described above, such that a total of five mock communities of varying compositions (Table S1) were prepared and sequenced on an Illumina MiSeq v3 at the University of Washington’s Center for Environmental Genomics. Sequencing data for the mock communities was processed using the same bioinformatics pipeline as for sample data.

Mock community and sample data were then used to run the Shelton et al. (2023) quantitative model in R v4.1.3. We ran the model using three chains, with 5000 iterations per chain, and the default tree length of 12. The estimated median proportion of DNA attributed to each calibrated taxa, for every crab included in the analysis, is reported in the Supplementary Material (Table S6). We subset the data by model run according to the total number of PCR cycles used the amplify / index each sample (i.e., replicate); for three crabs which were re-processed at two different total PCR cycles, we used only the data from the replicates run at the lowest total PCR cycles.

The resulting estimated proportions of prey DNA were then used to produce an “average” crab diet for each site type, by fitting a zero-inflated Dirichlet distribution with the R package zoid (Jensen et al., 2022; Ward et al., 2022). The zoid input data consisted of a matrix of estimated read counts for each calibrated prey species for each crab; each set of observations was treated as having been drawn from a multinomial distribution for which the probabilities of the *K* classes were the mean proportions of DNA per prey taxa estimated by the quantitative model, and the size *N* was the average read depth across technical replicates. Because of the nature of the data (high variability between individuals, with non-zero observations either very large or very small numbers), the likelihood-based optimizer used by ‘zoid’ had difficulty finding suitable initial values. We solved this by dividing all read depths in the input matrix by 100, effectively reducing our sample size while facilitating model fitting. This transformation is conservative, in the sense that it does not alter the mean model estimates (i.e., species compositions), but does somewhat inflate confidence intervals. We fit separate models with the *fit_zoid* function, using four chains with 10,000 iterations per chain, for crabs from clam bed sites and crabs from slough sites.

## Results

We dissected a total of 121 green crabs trapped at our four sampling sites (Fig 1, Table 1), 114 of which had enough stomach contents to sample; we successfully extracted, amplified, and sequenced DNA from 86 of these stomach content samples, 62 of which contained DNA from putative prey items (51 male / 9 female). Crab stomachs in which we detected prey DNA most commonly contained one to three distinct taxa (Fig. S2).

### Diet composition

We identified a total of 56 unique prey taxa across all crabs sequenced. Arthropoda was the most frequently observed phylum across all sampling sites and months, with 32 crab stomachs (52%) containing arthropod prey. Green crabs at the Oysterville clam bed site had the greatest α diversity (n=39 unique prey taxa; Table 1); this was also true when each site’s prey α diversity was divided by sample size to account for the fact that α diversity tended to scale with the number of crabs sequenced (Fig. S3). Adjusting for sample size, the Nahcotta (n=9 taxa; 1.5 average per crab) and Stackpole (n=16 taxa; 1.1 average per crab) slough sites had the second and third highest prey α diversity.

The single most common prey item was the native hairy shore crab (*Hemigrapsus oregonensis*), which was present in the stomach contents of crabs from all four sites (Fig 2, Table 2). After the hairy shore crab, the sand or shrimp (*Crangon franciscorum*), a gammarid amphipod (*Ampithoe valida*), and a family of eelgrasses/surfgrasses (*Zosteraceae*) were the most commonly identified taxa across all four sites (Fig 2, Table 2). Across the two clam bed sites, the Pacific staghorn sculpin (*Leptocottus armatus*), an invasive gammarid amphipod (*Gradidierella japonica*), and a genus of red algae (*Neoporphyra* sp.) were equally common, but were absent from stomach contents of crabs trapped at the two slough sites. Instead, green crabs trapped at the two slough sites most commonly consumed brown (*Fucus* sp.; *Leathesia marina*) and green (*Ulva compressa*) algae, a nereid worm (*Hediste diadroma*), and the soft-shell clam (*Mya arenaria*). Of these, only *M. arenaria* and *H. diadroma* were also present in the stomachs of crabs trapped on clam beds. Notably, despite trapping over half of the analyzed green crabs on Manila clam aquaculture beds, we found only one green crab with Manila clam DNA in its stomach contents (trapped in September at the Oysterville site).

**Figure 2.**
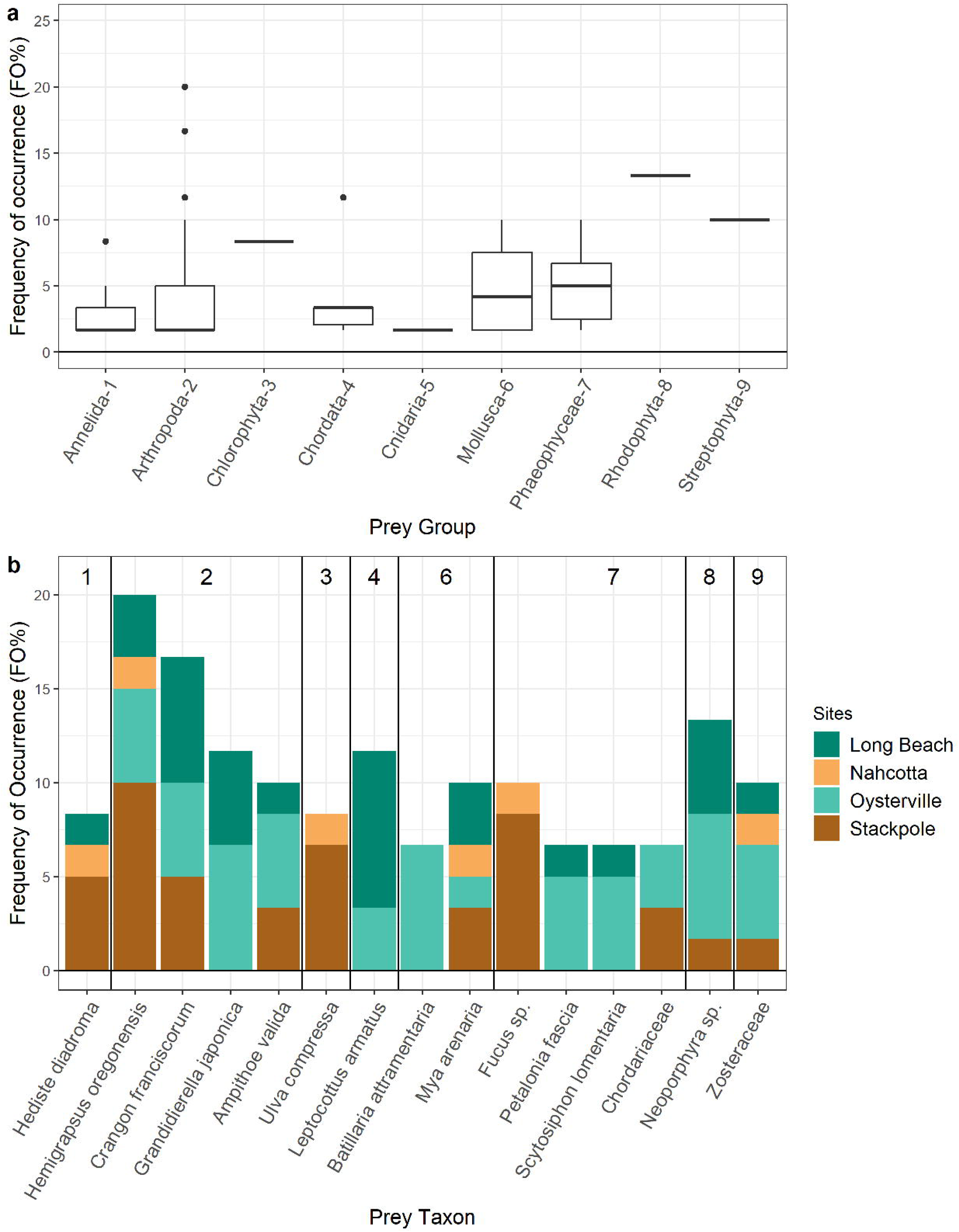
Frequency of prey occurrence, summarized (a) for major prey groups (distribution of FO% for individual taxa within each group, across all sites), and (b) for taxa which were present in four or more crab across all sites (total FO% broken down by site). Prey group identification numbers from (a) are annotated at the top of (b).

**Table 2.**
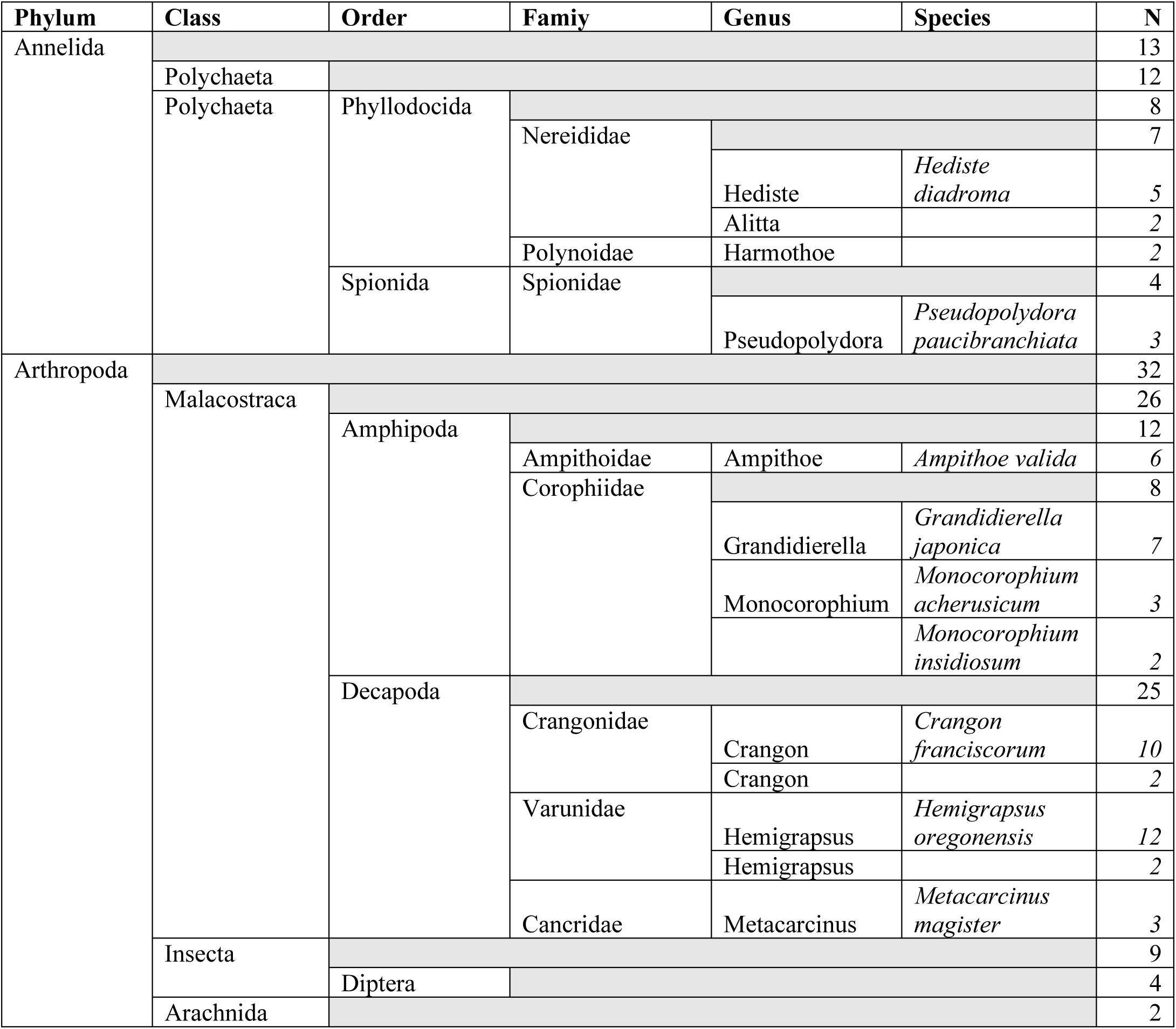
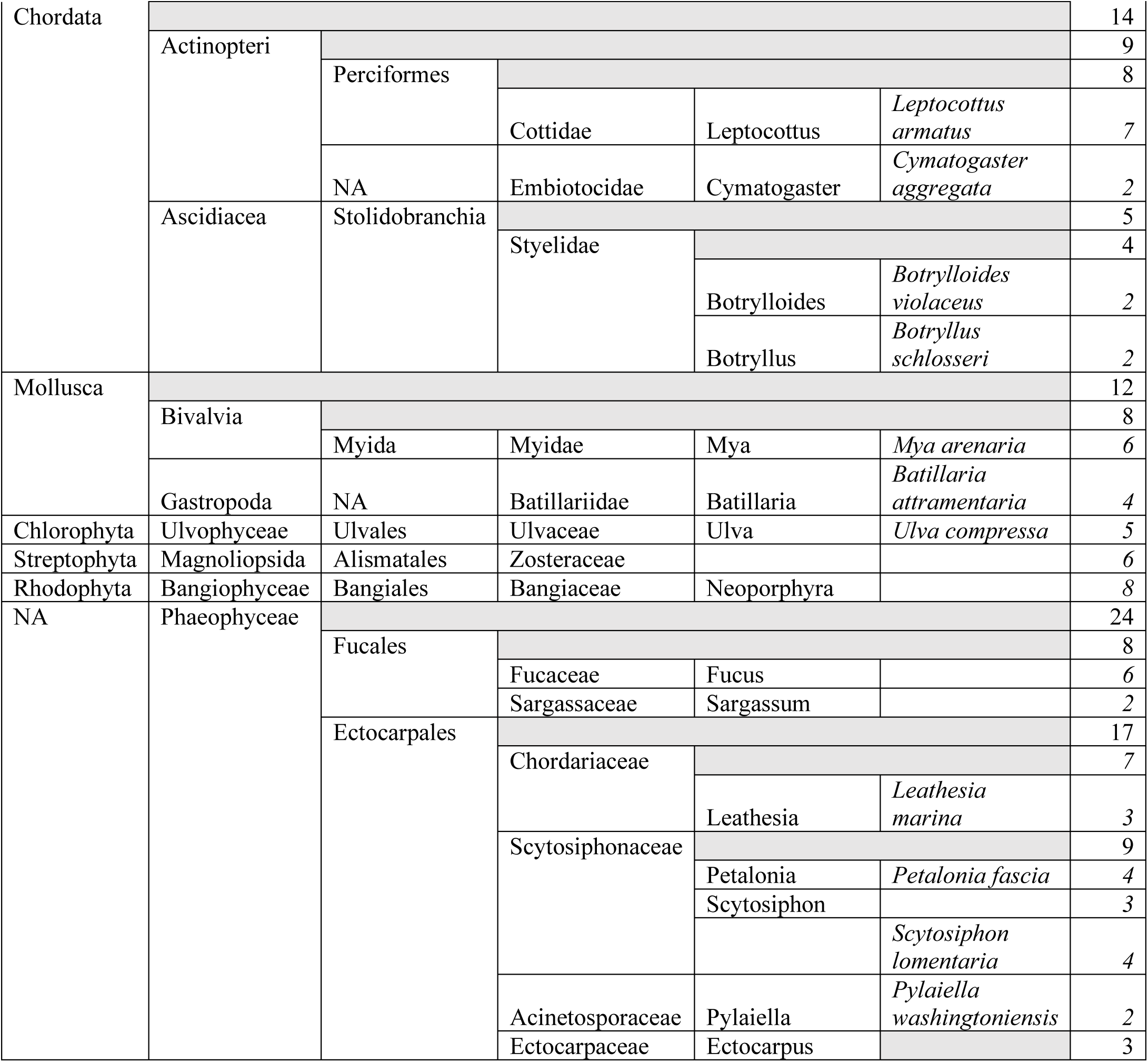
Prey items that occurred in two or more European green crab from any site, to their finest taxonomic classification, and the total number of crab in which that prey taxon was identified. Italicized numbers following gray shading represent totals for the given taxonomic level, summed across more specific taxonomies (including those not shown here, when n=1).

The traps used to collect green crabs for this study also captured bycatch species, several of which were identified in green crab stomach contents. All green crab with Dungeness crab in the stomach contents co-occurred with Dungeness crab in traps (Table S2). However, not all detections of bycatch species in stomach contents were associated with the co-occurrence of that species in the trap; for example, only one crab of the six identified as consuming Pacific staghorn sculpin (*Leptocottus armatus)*, four out of the twelve that consumed the hairy shore crab (*H. oregonensis)*, and none that consumed an invasive mud snail (*Batillaria attramentaria)* or the saddleback gunnel (*Pholis gunnellus)*, co-occurred with those species in the traps.

### Observed diet differences at aquaculture v. natural slough sites

Patterns in β diversity revealed that green crabs from sites of the same type (aquaculture, natural slough) had more shared prey species than green crabs from different site types. β diversity was lowest when comparing sites of the same type (e.g., between the two slough sampling sites, Nahcotta and Stackpole), and highest when comparing diet across site types (Table 3). The greatest difference in diet composition was observed between the Stackpole slough site and the Oysterville clam bed site (0.84; Table 3). In a Principal Coordinates Analysis (PCoA) using presence/absence data and the Jaccard distance, the two clam bed sites and two slough sites were clustered more closely to each other in coordinate space than they were to sites of the opposite type (Fig. 3). Further, while there were several strong outlier samples in the PCoA constructed with the eDNA index and the Bray-Curtis distance, the general pattern of similarity between sites of each type (clam bed vs. slough) remained.

**Figure 3.**
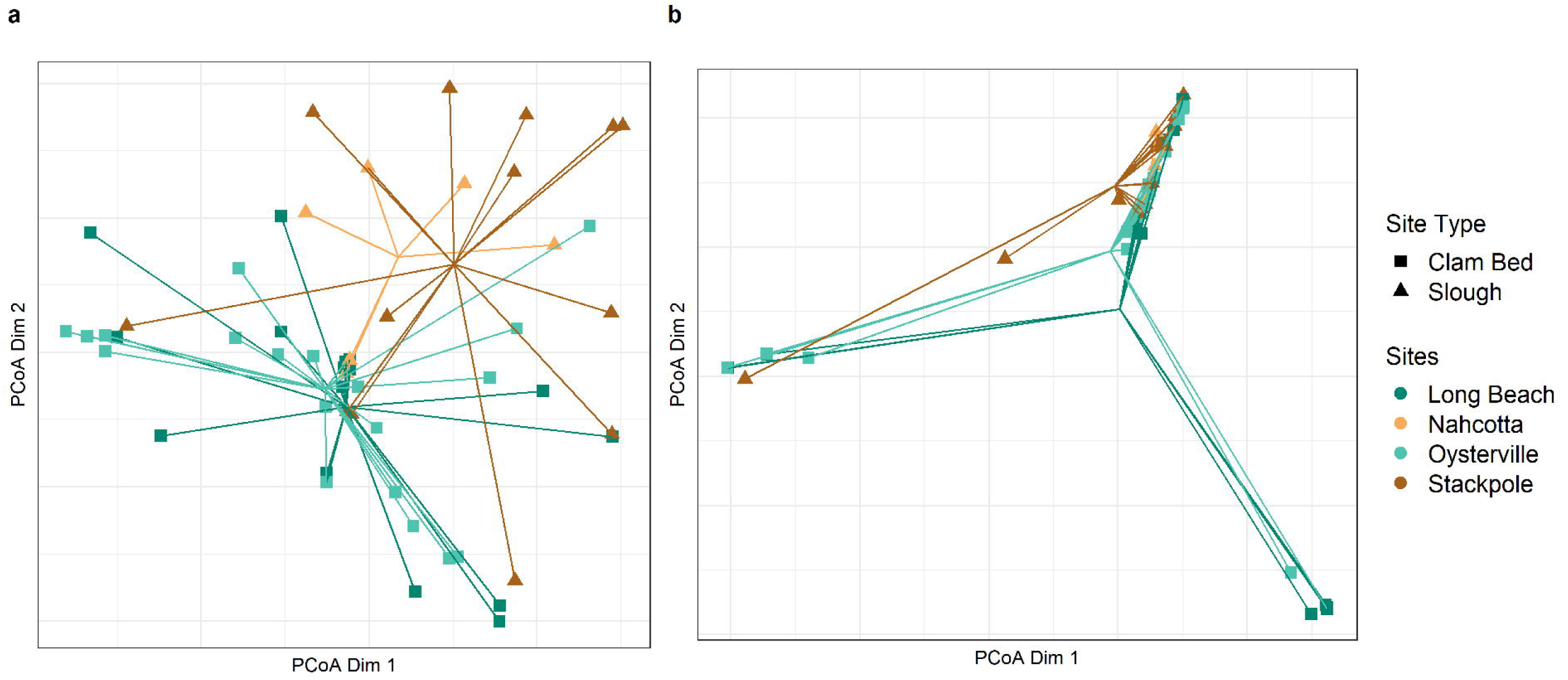
Principal Coordinate Analysis constructed using (a) a presence/absence matrix and the Jaccard dissimilarity index, and (b) the eDNA index and the Bray-Curtis distance.

**Table 3.**
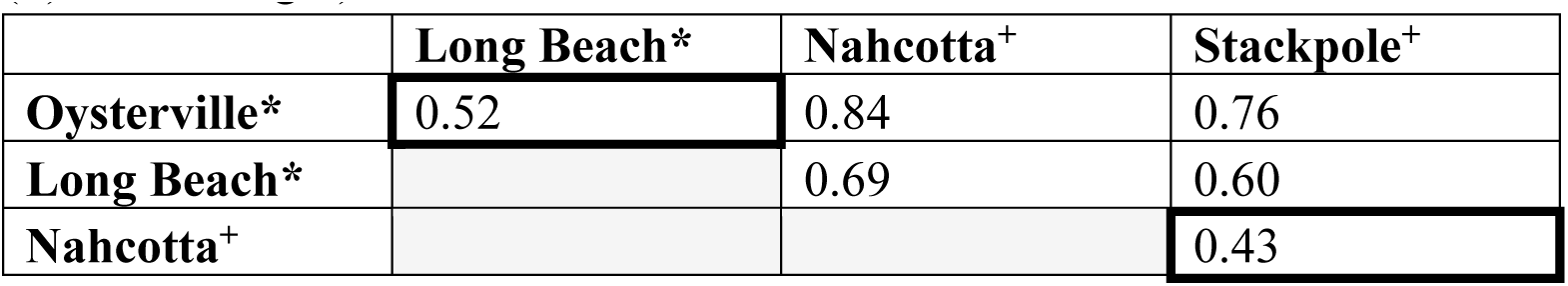
β diversity in prey items between the sampling sites. Darker borders outline the β diversities between the two clam bed sites (marked (*); top left) and the two slough sites (marked (+); bottom right).

Diet composition varied significantly between aquaculture versus natural slough sites. A PERMANOVA, run first with presence/absence data and then with the eDNA index values, found significant differences in diet composition between two or more of the sampling sites (p<0.05; Table S3). Post-hoc pairwise comparisons (Table S5) identified significant differences in diet composition between the crabs trapped at Stackpole (natural slough site) and those trapped at Long Beach (clam bed site; p < 0.05 for both data sets), and between Stackpole and Oysterville crabs (clam bed site; p < 0.05 for presence/absence data set alone). All other pairwise comparisons were non-significant. Our PERMDISPs returned non-significant results (Table S4), verifying that the significant PERMANOVAs were not due to differences in the dispersion of samples around the centroid of the Principal Coordinates Analysis (PCoA; Fig. 3).

### True abundance of key prey items in stomach content samples

To quantitatively compare how different prey of interest contributed to green crab diets, we calculated true prey DNA abundance with our mock community data and the Shelton et al. (2023) quantitative model. This mechanistic model calibrates sample data relative to a single reference species (Shelton et al., 2023); a higher amplification efficiency means that a given species may be over-represented in the DNA metabarcoding data compared to the reference. With Manila clam as the reference species, green crab had the highest relative amplification efficiency for our BF3/BR2 primer set, followed by the shrimp *C. franciscorum,* and Dungeness crab (Table 4). The native eelgrass *Z. marina* was the only species with a lower amplification efficiency than Manila clam (Table 4).

**Table 4.** Estimates of amplification efficiencies (α) for all species in the mock communities. Amplification efficiencies are relative to the Manila clam. ^+^ Not present in sample data, α estimated from separate run of the quantitative model. The α parameter captures the log-ratio of amplification efficiency for a species relative to that of the reference species (here, Manila clam; see Shelton et al. 2023 for full description of α).

| Species (Common Name) | $\alpha$ |
| --- | --- |
| *Ruditapes philippinarum* (Manila clam) | 0 |
| *Zostera marina* (eelgrass) | -0.0098<sup>+</sup> |
| *Cottus asper* (prickly sculpin) | 0.0360<sup>+</sup> |
| *Hemigrapsus oregonensis* (hairy shore crab) | 0.0402 |
| *Leptocottus armatus* (Pacific staghorn sculpin) | 0.0478 |
| *Batillaria attramentaria* (invasive mud snail) | 0.0607 |
| *Mya arenaria* (soft-shell clam) | 0.0619 |
| *Cymatogaster aggregata* (surf perch) | 0.0687 |
| *Cancer magister* (Dungeness crab) | 0.0817 |
| *Crangon franciscorum* (sand shrimp) | 0.0859 |
| *Carcinus maenas* (European green crab) | 0.0896 |

Of the 62 green crabs with prey taxa in their stomach contents, 34 contained DNA from one or more of the calibrated prey species (i.e., those for which we had amplification efficiencies from the quantitative model). We did not detect the prickly sculpin (*C. asper*) in green crab stomach contents, and although the family *Zosteraceae* was present in stomach contents, it would be inappropriate to apply our calibration derived from *Z. marina* to an identification that may represent a mixture of different eelgrass and surf grass species (e.g., *Z. japonica* or *Phyllospadix torreyi*, both of which are present in Willapa Bay) . On average, the eight calibrated prey species that were also in green crab stomach contents made up less than 50% of the unique prey taxa found in each crab stomach, but there was wide variation in this statistic among crabs: some crabs collected from the two clam bed sites only had calibrated prey detected in their stomach contents (Fig. 4b). Among crabs collected on clam beds, the calibrated prey represented, on average, almost 100% of all sequence reads from prey taxa, whereas for crabs collected at the slough sites, 50% of reads on average came from non-calibrated prey taxa (Fig. 4b).

**Figure 4.**
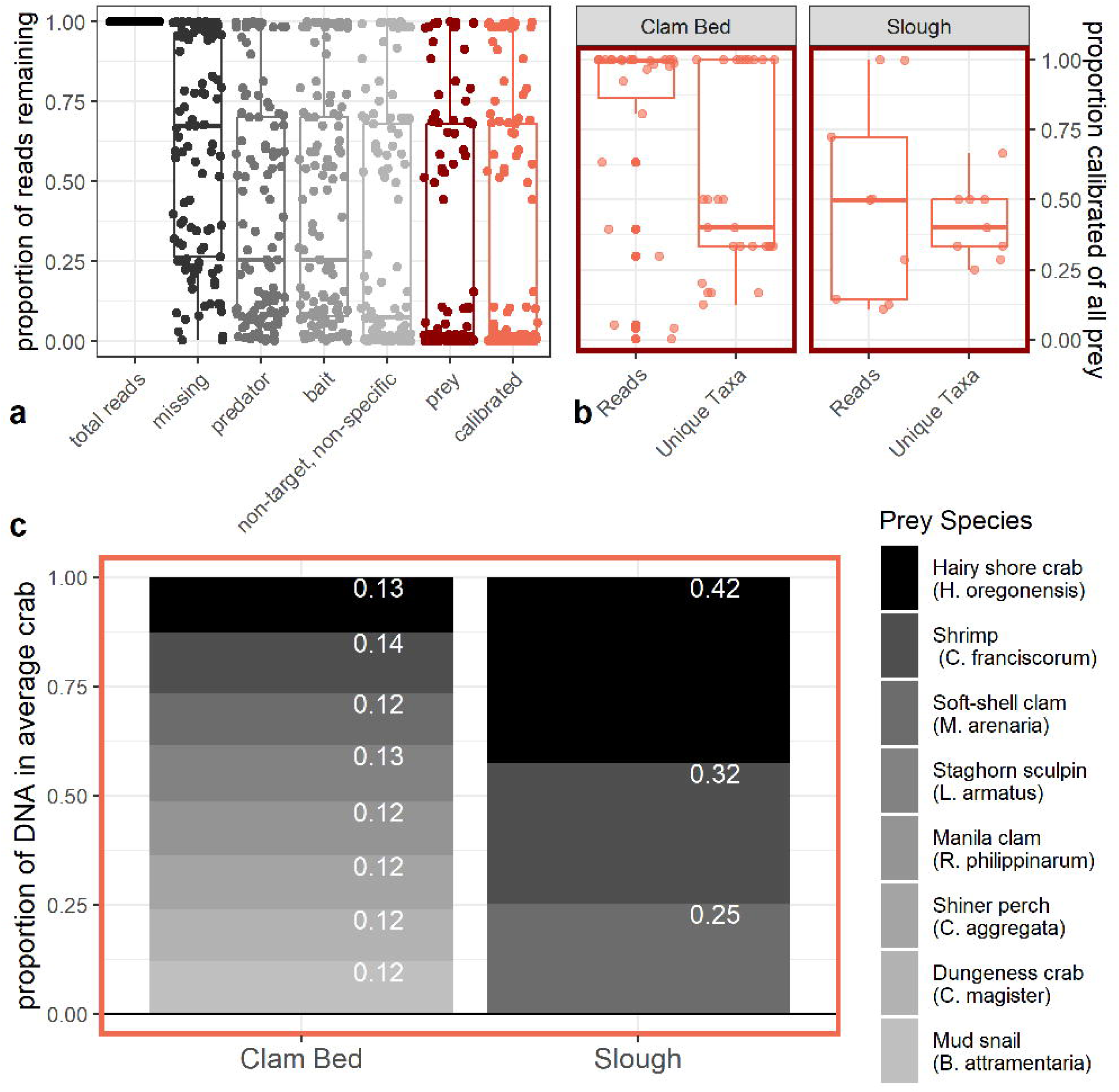
The diet of an “average” green crab which contains the eight calibrated prey species, from each site type **(c)**. Proportions for each prey species are provided in white text in the appropriate block, and represent the mean estimate from the fitted zero-inflated Dirichlet distribution. To put into perspective what subset of all sequencing data is represented by the eight calibrated prey species, we also provide **(a)** the proportion of sequencing reads remaining after each filtering step, out of the total matched to the blast database (“total reads”), and **(b)** the proportion of reads belonging to all prey taxa, and proportion of all unique prey taxa, captured by the eight calibrated prey species (pink). Confidence intervals associated with the mean values in **(c)** can be found in Fig. S6.

As in the full dataset, the hairy shore crab (*H. oregonensis*) was the most frequently occurring prey item of those calibrated with the Shelton et al. (2023) model (n = 11 / 34 green crabs), followed by the sand shrimp (*C. franciscorum;* n = 9 / 34 green crabs). When it was detected in a green crab stomach, the sand shrimp also contributed the highest average proportion of DNA out of all calibrated prey species, followed by the Pacific staghorn sculpin and the Manila clam(although there was variation in DNA proportions across crab, and in sample sizes across prey species; Fig. S4; Table S8).

The sand shrimp (*C. franciscorum*) and the Pacific staghorn sculpin (*L. armatus*) were similarly ranked in the calibrated DNA data and the uncalibrated sequencing read data (highest average proportion of sequencing reads and average proportion of DNA out of the calibrated species, when detected in the stomach contents; Table S8). However, the Manila clam increased in rank from the uncalibrated (rank = 8/8) to the calibrated (rank = 3/8) data set; and in contrast, the ranked abundance of the soft-shell clam (*M. arenaria*) decreased from the uncalibrated (4/8) to the calibrated (8/8) data set due to its high relative amplification efficiency (Fig S4; Table S8).

### Average green crab diet at aquaculture v. natural slough sites

Modeled average diet composition at the clam bed sites showed nearly even contributions to the diet from each of the eight calibrated prey species (Fig. 4c; Table S7), with the sand shrimp (*C. franciscorum*) contributing a slightly higher mean proportion of DNA than the other species (0.14 of total DNA; 95% confidence interval (0.12,0.18)). In contrast to the clam bed sites, only three of the eight calibrated prey were identified in green crabs collected at the two slough sites. The majority of the DNA in an “average” crab stomach from the slough sites was estimated to come from the native hairy shore crab (*H. oregonensis*; 0.42 of total DNA; 95% confidence interval (0.31,0.54)), followed by the sand shrimp (*C. franciscorum*; 0.32 of total DNA; 95% confidence interval (0.32,0.24)), and the soft-shell clam (*M. arenaria;* 0.25 of total DNA; 95% confidence interval (0.16,0.35); Fig. 4c; Table S7).

## Discussion

Our research provides the first detailed diet information for the invasive European green crab in the northeastern Pacific. These data are particularly valuable in Washington State, where an “exponential” increase in green crab populations prompted an emergency proclamation by the state governor (WA Governor’s Emergency Proclamation No. 22-02 Jan 19 2022). We found that green crab in Willapa Bay Washington have a generalist, omnivorous diet that can be highly variable across individuals and locations. Even the most common taxon identified in the stomach contents, the hairy shore crab (*H. oregonensis*), had a frequency of occurrence of only ∼ 20%. The role of green crab as a generalist predator in Willapa Bay agrees with existing research from other locations where the species is invasive (Cordone et al., 2022; Le Roux et al., 1990), and within its native range (Ansell et al., 1999; Baeta et al., 2006). However, our use of DNA metabarcoding allowed for a greater number of prey taxa to be identified from a smaller number of crabs (Ansell et al., 1999) and at a higher taxonomic resolution (Le Roux et al., 1990) than prior research using visual diet analysis.

We also distinguished statistically significant differences in stomach contents between sampling sites; green crabs trapped in the natural slough at the Stackpole site had significantly different diet compositions than those trapped on active Manila clam aquaculture beds at Long Beach and Oysterville. Although we did not concurrently examine ecological communities at sampling sites, we suspect this is reflective of different community compositions on Manila clam beds and uncultured slough sites, rather than indicative of differing prey selectivity among green crabs at these sites. Previous work on green crab diet on the Patagonian coast found strong similarities between their most frequently occurring prey items and surveys of the local ecological community during separate benthic monitoring (Cordone et al. 2022).

### Congruence with existing research

The only other study of green crab diet using DNA metabarcoding (diet DNA or dDNA) was conducted on the Patagonian Atlantic coast (Cordone et al., 2022). As in the present study, Cordone et al. (2022) reported Arthropoda as the most frequently identified phylum in green crab stomach contents, and noted the diversity of algal taxa consumed, particularly the class *Phaeophyceae,* which had not been documented in prior visual diet analyses. Cordone et al. (2022) suggested that algae in the stomach contents most likely represents incidental ingestion, based on their concurrent findings from stable isotope analysis that the energy pathway for macroalgae was less important than that for phytoplankton. However, this does not necessarily exclude the potential for green crab to have negative impacts on macroalgal abundance and diversity.

In contrast to Cordone et al. (2022), bony fishes had relatively high frequencies of occurrence in our dataset, particularly the Pacific staghorn sculpin (*Leptocottus armatus*). Other studies have found juvenile fish in green crab stomach contents (Ansell et al., 1999; Taylor, 2005), with increasing crab sizes associated with more frequent fish consumption (Taylor, 2005). Our sampling process, in which we selectively analyzed larger crabs, may therefore explain the greater relative importance of fish species in our dataset than from Cordone et al. 2022 (14.5% FO, compared to 5.71% FO). Yet where Taylor (2005) estimated that green crab predation has a limited impact on winter flounder (*Pseudopleuronectes americanus*) populations in Connecticut, there may be greater potential for population-level impacts in Willapa Bay, given the frequency of *L. armatus* consumption. Moreover, the authors have personally observed Pacific staghorn sculpin aggregating near actively foraging crab, and believe it is possible sculpin may be opportunistically captured and consumed while crabs are foraging for other prey.

### Implications for Willapa Bay ecosystems

The present work was initiated because of concern for Manila clam aquaculture in Willapa Bay, where unexplained declines in production have occurred. Previous work in California has shown green crab to be effective predators on cultured Manila clams (Grosholz et al., 2001), and recent increases in the abundance of green crab in coastal Washington waters have been hypothesized to contribute to the observed clam production declines. However, the present study does not provide substantial support for this speculation; only one sampled green crab showed evidence of consuming Manila clam. Notably, this observation occurred in a clam aquaculture area (Oysterville). Both clam beds sampled were under active culture with a wide size range of clams present, and no netting or bags were used to protect clams from predatory crabs (as is often done in other aquaculture settings).

It is possible that green crabs could be exerting higher predation rates on clam beds outside of our summer sampling period; though consumption and activity rates of green crabs drop in colder temperatures (Aagaard et al., 1995; Elner, 1980), their migration to lower elevations often means that larger relative abundances are observed on Willapa Bay Manila clam beds in winter months. Additionally, size class matches between predator and prey remain an open question for this system. We selected large crabs for diet analysis, but small Manila clams may only be profitable for smaller size classes of green crabs; thus impacts by green crabs may be stage and context dependent, and most severe in habitats where smaller green crabs are found on commercial clam beds. As a result, while Manila clams were an extremely rare diet item in the green crabs we analyzed with the BF3/BR2 primer set, more extensive sampling throughout the year and across smaller size classes may yield different results. Manila clam did have a lower relative amplification efficiency of all but one of the species calibrated in the Shelton et al. (2023) model, particularly compared to green crab itself; yet we had no missed detections and adequate read depths of Manila clam in mock communities, which included the species’ DNA at low relative proportions to green crab and other species (e.g., Community D; Table S1). Given the strong signal of Manila clam in mock communities, we have confidence that if Manila clam had been recently consumed by the green crab we analyzed, we would have detected it in the stomach contents.

In contrast to Manila clam, we detected soft-shell clams (*Mya arenaria*) in the diet of multiple green crabs across several sites. Green crabs have been implicated in the collapse of a soft-shell clam (*M. arenaria*) fishery in Maine in the mid-twentieth century, when clam abundance declined by 50% in a 4-year period (Glude, 1955). Green crabs have been shown to preferentially consume soft-shell clams in some studies (Miron et al., 2005; Pickering & Quijón, 2011) and substantial effort has been allocated toward strategies to reduce predation in the northeastern Atlantic (Floyd & Williams, 2004; Tan & Beal, 2015). Soft-shell clams are non-indigenous to the northeastern Pacific, but have established large populations across Willapa Bay since their initial introduction prior to 1880; soft-shell clams may therefore represent an important food resource for green crab in this estuary.

Although we did detect three cases of green crab consumption of Dungeness crab (*Cancer magister;* at Long Beach and Oysterville, in May), Dungeness crabs were recorded as bycatch in the same traps corresponding to these three green crabs. It is therefore possible that consumption of Dungeness crab (*C. magister*) was facilitated by trapping, and predation would not otherwise have been observed. Notably, however, we did not find physical evidence of predation occurring within the crab traps (i.e., carapace fragments or other debris). We also limited soak times and retrieved traps promptly to reduce the likelihood of consumption occurring within the traps. Nevertheless, hand capture could reduce the likelihood of diet observations being confounded by predator-prey interactions attributable to sampling technique.

Other arthropods observed in stomach contents from this study, while not a focus of commercial aquaculture and fisheries, play an important ecosystem role in Pacific Northwest estuaries. In the present study, a genus of sand shrimps (*Crangon spp*.) occurred in more than 20% of all samples –primarily as *Crangon Franciscorum,* when identified to the species level. Our quantitative modeling results further show that when the sand shrimp (*C. franciscorum*) was consumed by a green crab, the species comprised, on average, a high proportion of DNA in the stomach content sample (compared to other calibrated prey). This particular species (*C. franciscorum*) is the predominant sand shrimp in Willapa Bay and an important predator, feeding on amphipods, bivalves, and small or juvenile fish (Wahle, 1985). The species is also an important food resource for Dungeness crab (*C. magister;* Stevens et al., 1982), sturgeon, and other fishes. Their sister taxon, *C. crangon*, is known to be an important diet component in the green crab’s native European range (Baeta et al., 2006) in all seasons. We also found that a diversity of amphipods occurred in 29% of the sampled green crabs across all sites. As a taxonomic group, amphipods play a critical role in nutrient cycling and other ecosystem functions and are important prey to a wide variety of invertebrates, fishes and birds. Green crab is known to frequently consume amphipods in both the native (Bleile & Thieltges, 2021) and invasive range (Donahue et al., 2009; Le Roux et al., 1990), and significantly reduce their densities in enclosure experiments (Grosholz & Ruiz, 1995).

Prior work has also demonstrated the negative impacts of green crab on native shore crabs, one species of which (*H. oregonensis*) was the most frequently occurring taxon in our dataset of green crab diets. In the northeastern Pacific, impacts to native shore crab have been extensively documented in central California (Grosholz et al., 2000; de Rivera et al., 2011). In particular, native shore crab numbers declined and distribution shifted into the high intertidal zone as green crab became more abundant. Moreover, the average size of the native shore crab declined at the same time (de Rivera et al., 2011). Given the evidence of predation on this species observed in Willapa Bay, as well as the well-documented overlap in habitat use, it seems likely that similar population-level impacts could also occur in Washington State estuaries.

One difficulty in interpreting diet data is that we cannot determine whether the presence of a species in the stomach contents is the result of direct or intentional predation. For instance, the detection of some taxa might occur because they were attached to (epibiont), inside of (themselves prey of green crab prey), or otherwise associated with items green crabs targeted as prey. For example, Ansell et al. (1999) dissected a complete juvenile fish in a green crab stomach, and found that the fish had previously consumed copepods, ostracods, and an amphipod. We detected all three of these taxonomic groups in our green crab stomach contents. However, we developed our post-BLAST filtering process to select for taxa which were most likely in the size range of true prey for the crabs we sampled, and previous work with terrestrial arthropods suggests that using amplicons greater than 200bp in length reduces the likelihood of detecting DNA from scavenged prey (Greenstone et al., 2014), which may have levels of DNA degradation similar to items representing secondary predation. Incidental ingestion might also account for some detections in our study, our identification of *Zosteraceae* (a family of surfgrasses and eelgrasses) DNA in particular. While green crabs are known to eat eelgrass, both as seeds and blades in experimental (Infantes et al., 2016) and field enclosures (Howard et al., 2019), respectively, several authors have suggested that consumption may occur incidentally during other foraging activities (Davis et al., 1998; Garbary et al., 2014; Malyshev & Quijón, 2011). As mentioned above, algae may also be accidentally consumed during other foraging activities; generally this category represents a small component of the diet relative to known preferred prey like molluscs (Baeta et al., 2006; Cordone et al., 2022; Le Roux et al., 1990).

Although beyond the purpose of our study, which focused on green crab predation on other organisms, one key aspect of crab diet that cannot be captured using DNA metabarcoding is cannibalism. Cannibalism is common in decapod crustaceans, and has been shown to cause high density-dependent mortality in green crab (Moksnes 2004, Grosholz et al., 2021). However, DNA metabarcoding cannot distinguish between individuals of the same species, and so all green crab DNA in our data was assumed to originate from the predator individual. Future research with the goal of understanding cannibalism in Pacific Northwest green crab populations could employ other molecular techniques to do so, such as genotyping of microsatellite loci, which has revealed cannibalism in invasive lionfish populations (Dahl et al., 2018).

### Moving toward quantitative dDNA for invasive species impact assessments

Calibrating DNA metabarcoding data to account for species-specific primer amplification bias provided key insights into our own data set, and contributes important information for future studies of green crab predation. The rank abundance of prey species’ contributions to green crab stomach contents changed after we calibrated sequence read abundances; both the hairy shore crab (*H. oregonensis*) and the Manila clam increased in rank abundance in the calibrated DNA data compared to the sequence read data, whereas the soft-shell clam *M. arenaria* decreased in rank abundance (Table S8). Our calibrated data set revealed that out of the eight calibrated species, the sand shrimp (*C. franciscorum*), the Pacific staghorn sculpin (*L. armatus*), and the Manila clam contributed the highest average amount of DNA when present in the stomach contents. By applying the calibrated data to model an average green crab stomach sample, our research also emphasized differences in relative abundance of eight key prey items between aquaculture versus natural slough sites; at slough sites, the native hairy shore crab (*H. oregonensis*) had a higher average abundance in the stomach content DNA than the soft-shell clam (*M. arenaria*), whereas at the clam bed sites, we did not observe clear differences in average DNA abundance between the eight calibrated prey species.

In generating the calibrated data set, we also revealed that green crab itself had the highest PCR amplification efficiency of all the species included in the mock communities; this was not entirely unexpected, as the BF3/BR2 primer pair was initially developed to study benthic invertebrates (Wangensteen et al., 2018), but it does suggest that exploring alternative primer sets may help future work minimize the loss of sequencing reads to predator DNA. With mock communities and the Shelton et al. (2023) model, future studies can use different primers and still be comparable to this work. The calibration of additional prey taxa that were particularly prevalent in the diet of crabs collected for this study – such as the red algae genus *Neoporphyra*, and the invasive amphipod *Grandidierella japonica* – will be important for assessing the broader ecological impact of green crabs in the Pacific Northwest, and will help put into perspective the relative contribution of the prey calibrated for this study in DNA extracts from green crab stomachs.

A major target for applying diet DNA to invasive species research and management is population-level impact analysis, and the potential for DNA metabarcoding to provide quantitative consumption data for modeling methods like bioenergetics. As the first published application of the Shelton et al. (2023) model to stomach content data, our research brings the study of invasive European green crab one step closer to this goal; however, the proportion of observed DNA contributed by each prey species (the “true DNA abundance” output from the Shelton et al. 2023 model) is not yet translatable to the proportion of consumed biomass contributed by each prey species (input for bioenergetic modeling). Without accounting for changes in detectability as prey DNA degrades during digestion, quantitative interpretations of dDNA may overemphasize predation impact on prey with lower DNA digestion rates, and, conversely, underestimate predation impact on prey with higher DNA digestion rates (Greenstone et al., 2014). A meta-analysis of spiders found that accounting for differential prey detectability could change dietary proportions of a given spider prey item by up to ∼10% (Uiterwaal & DeLong, 2020). Experimental feeding trials that explore how the detectability of prey DNA changes with time since ingestion, often quantified using the genetic or detectability half-life (Dick et al., 2023; Greenstone et al., 2014), will be key to additional applications of our research and the broader integration of the Shelton et al. (2023) model into invasive species impact analysis. Currently, there are no published feeding experiments to quantify prey detectability half-lives for green crab. Since detectability half-life varies across prey species, feeding experiments to determine prey half-lives can be particularly prohibitive for generalist opportunists like green crab; however, a recent meta-analysis for terrestrial arthropods suggests that there may be consistent and predictable effects of prey traits and environmental conditions on detectability half-lives that make it possible to forego exhaustive feeding trials leveraging meta-analyses and quantitative models (Uiterwaal & DeLong, 2020).

## Conclusions

- Our research suggests that several ecologically important native species, including the hairy shore crab (*H. oregonensis*), the Pacific staghorn sculpin (*L. armatus*), and the sand shrimp (*C. franciscorum*), are strong candidates for experiencing population-level impacts from increasing green crab populations in Washington State estuaries. Ongoing efforts to monitor these species, as well as other community-level shifts in trophic structure and vegetation cover, will be critical to the early detection of negative impacts. Monitoring efforts, and any further diet analyses, should be completed alongside continued investment in efforts to reduce the spread and growth of invasive green crab populations.
- We observed significant differences in green crab stomach contents between the natural intertidal slough sites and the Manila clam aquaculture sites, although the extent to which this reflects the presence of ecologically distinct communities at the two site types, as opposed to differing prey selectivity, requires further investigation.
- To operationalize our DNA metabarcoding data for predation impact analysis, future research must include laboratory feeding trials to quantify the detectability half-lives of the prey items for which we provide amplification efficiencies. More broadly, conducting this combination of complementary research will further integrate DNA-based diet analysis into invasive species research and management.

## Acknowledgements

Funding was provided by the Washington Department of Fish and Wildlife (WDFW No.21-18297) as part of the Willapa Bay Oyster Reserves Grant Program (RFP No. 21-0009). Mary Fisher was also supported by the NSF’s Graduate Research Fellowship Program (Grant DGE-1762114) while conducting this work. Green crab collection was conducted by Washington Sea Grant Crab Team and the Willapa Brays Harbor Oyster Growers’ Association under WDFW Aquatic Invasive Species Permit EGC.001-21.20. We would like to thank Friday Harbor Laboratories and the University of Washington’s Burke Museum Fish Collection for their assistance in obtaining individuals and tissue samples used for mock communities. Thank you also to Heckes’ Clam Company, Taylor Shellfish, and Kim Patten for providing property access for sample collection. Bettina Thalinger, Georgina Cordone, Eily Allen, Erin D’Agnese, Megan Shaffer, and Maya Garber-Yonts shared invaluable lab and bioinformatic knowledge and assistance, and Eric Ward provided help troubleshooting the R package zoid.

