## Supplemental Tables, Figures, Text for "Invasive European green crab (*Carcinus maenas*) predation revealed with quantitative DNA metabarcoding"

### S1 Text. Trapping Protocol

Galvanized steel minnow traps (Gee's G-40), and square Fukui fish traps were baited with Pacific mackerel (*Scomber japonicus*) and set for the nighttime high tide. At clam bed sites, traps were arrayed in 5 horizontal transects of 6 traps each, alternating trap type. Traps within each horizontal transect, and each of the five horizontal transects, were separated by approximately 20m, such that the array formed a checkerboard grid. In natural sloughs, 10 traps of each type were set in the middle of the slough following the main channel, starting at the highest elevation at which halophyte vegetation was apparent on the shore, and extending down channel, alternating trap type, again with each trap separated by 20m. On trap retrieval, all organisms in each trap were recorded to species, and a subset of all crab species were measured (carapace width) to the nearest mm. Native species were released, with the exception of Dungeness crabs (*Cancer magister*) which were retained for another project.

**S1 Table. Mock community composition.**

| Label | Descriptor | Species | Relative proportion | Total DNA (ng) |
|---|---|---|---|---|
| A | All species, even proportions | *C. maenas, R. philippinarum, M. arenaria, C. magister, H. oregonensis, C. franciscorum, B. attramentaria, Z. marina, C. aggregata, L. armatus, C. asper* | 1 | 20 |
| B | No green crab, even proportions | *R. philippinarum, M. arenaria, C. magister, H. oregonensis, C. franciscorum, B. attramentaria, Z. marina, C. aggregata, L. armatus, C. asper* | 1 | 20 |
| C | No crab or shrimp, even proportions | *R. philippinarum, M. arenaria, B. attramentaria, Z. marina, C. aggregata, L. armatus, C. asper* | 1 | 20 |
| D | All species, skewed proportions | *C. maenas* | 3.3 | 66 |
| | | *R. philippinarum* | 0.88 | 17.60 |
| | | *M. arenaria* | 1.01 | 20.20 |
| | | *C. magister* | 0.30 | 6 |
| | | *H. oregonensis* | 0.45 | 9 |
| | | *C. franciscorum* | 0.77 | 15.40 |
| | | *B. attramentaria* | 0.74 | 14.80 |
| | | *Z. marina* | 0.98 | 19.60 |
| | | *C. aggregata* | 1.10 | 22 |
| | | *L. armatus* | 0.93 | 18.60 |
| | | *C. asper* | 0.95 | 19 |
| E | No green crab, skewed proportions | *R. philippinarum* | 1.60 | 32 |
| | | *M. arenaria* | 1.20 | 24 |
| | | *C. magister* | 0.30 | 6 |
| | | *H. oregonensis* | 0.45 | 9 |
| | | *C. franciscorum* | 1.50 | 30 |

| | | | | |
|---|---|---|---|---|
| | | *B. attramentaria* | 1.09 | 21.80 |
| | | *Z. marina* | 1.80 | 36 |
| | | *C. aggregata* | 1.70 | 34 |
| | | *L. armatus* | 1.40 | 28 |
| | | *C. asper* | 0.65 | 13 |

**S2 Table. Prey items found in green crab stomach contents that are also bycatch species.** We provide the number of crab with the prey item in its stomach, the number of those crab which also co-occurred with the given prey item in the trap, and the percent of crab that did *not* co-occur with the given prey item in the trap. The crab at Long Beach which had *Hemigrapsus sp.* identified from its stomach contents co-occurred with one *Hemigrapsus oregonensis* in the trap.

| Prey taxa | Site | N Crab | N Crab **with** co-occurrence | Percent Crab **without** co-occurrence |
|---|---|---|---|---|
| *Batillaria attramentaria* | Oysterville | 4 | 0 | 100 |
| *Hemigrapsus sp.* | Long Beach | 1 | 1 | 0 |
| *Hemigrapsus oregonensis* | Long Beach | 2 | 0 | 100 |
| *Hemigrapsus oregonensis* | Nahcotta | 1 | 0 | 100 |
| *Hemigrapsus oregonensis* | Oysterville | 3 | 3 | 0 |
| *Hemigrapsus oregonensis* | Stackpole | 6 | 1 | 83 |
| *Leptocottus armatus* | Long Beach | 4 | 1 | 75 |
| *Leptocottus armatus* | Oysterville | 2 | 0 | 100 |
| *Cancer magister* | Long Beach | 2 | 2 | 0 |
| *Cancer magister* | Oysterville | 1 | 1 | 0 |
| *Pholis gunnellus* | Oysterville | 1 | 0 | 100 |

**S3 Table. PERMANOVA results to compare diet composition between sampling sites**, based on (a) presence/absence, and (b) the eDNA index.

**(a)**

| | DF | Sum of Squares | R2 | F | Pr(>F) |
|---|---|---|---|---|---|
| site | 3 | 2.0161 | 0.07287 | 1.4671 | 0.004** |
| Residual | 56 | 25.6512 | 0.92713 | | |
| Total | 59 | 27.6672 | 1 | | |

**(b)**

| | DF | Sum of Squares | R2 | F | Pr(>F) |
|---|---|---|---|---|---|
| site | 3 | 1.9568 | 0.06958 | 1.396 | 0.011* |
| Residual | 56 | 26.1664 | 0.93042 | | |
| Total | 59 | 28.1232 | 1 | | |

**S4 Table. Evaluation of sample dispersion around centroids with the PERMDISP**, based on (a) presence/absence, and (b) the eDNA index.

**(a)**

| | **DF** | **Sum of Squares** | **R2** | **F** | **Pr(>F)** |
|---|---|---|---|---|---|
| group | 3 | 0.019514 | 0.09915 | 2.0544 | 0.124 |
| Residual | 56 | 0.177307 | 0.90085 | | |
| Total | 59 | 0.196821 | 1 | | |

**(b)**

| | | | | | |
|---|---|---|---|---|---|
| group | 3 | 0.006135 | 0.04849 | 0.9513 | 0.4 |
| Residual | 56 | 0.120378 | 0.95151 | | |
| Total | 59 | 0.126513 | 1 | | |

**S5 Table. Statistically significant permutational pairwise tests of diet composition between sites**, with ("corrected") and without Bonferroni correction applied to the p-value, based on (a) presence/absence, and (b) the eDNA index. Superscript indicates significance at the α=0.10 (.), 0.05 (*), and 0.01 (**) levels.

**(a)**

| | **DF** | **Sum of Squares** | **R2** | **F** | **P value** | **P value corrected** |
|---|---|---|---|---|---|---|
| *Oysterville v. Stackpole* | | | | | | |
| site | 1 | 0.81 | 0.05043 | 1.7524 | 0.001** | 0.006* |
| Residual | 33 | 15.253 | 0.94957 | | | |
| Total | 34 | 16.063 | 1 | | | |
| *Long Beach v. Stackpole* | | | | | | |
| site | 1 | 0.9093 | 0.05989 | 1.975 | 0.001** | 0.006* |
| Residual | 31 | 14.2727 | 0.94011 | | | |
| Total | 32 | 15.1821 | 1 | | | |

**(b)**

| | | | | | | |
|---|---|---|---|---|---|---|
| *Oysterville v. Stackpole* | | | | | | |
| site | 1 | 0.81 | 0.05043 | 1.7524 | 0.012* | 0.072 . |
| Residual | 33 | 15.253 | 0.94957 | | | |
| Total | 34 | 16.063 | 1 | | | |
| *Long Beach v. Stackpole* | | | | | | |
| site | 1 | 0.9093 | 0.05989 | 1.975 | 0.004** | 0.024* |
| Residual | 31 | 14.2727 | 0.94011 | | | |
| Total | 32 | 15.1821 | 1 | | | |
| *Oysterville v. Nahcotta* | | | | | | |
| site | 1 | 0.6086 | 0.04868 | 1.2792 | 0.098 . | 0.588 |
| Residual | 25 | 11.8937 | 0.95132 | | | |
| Total | 26 | 12.5022 | 1 | | | |

**S6 Table. Extension of Table 2, for all prey taxa.** See separate excel document.

**S7 Table. Mean and median estimated proportions of DNA contributed by calibrated prey species to an "average" crab diet, with confidence intervals.** Proportions are provided across all sites, and then for each individual site type (clam bed sites = Oysterville, Long Beach; slough sites = Nahcotta, Stackpole).

| Group | Species | Common name | Mean | Median | 95% CI |
|---|---|---|---|---|---|
| Clam bed | *Crangon franciscorum* | sand shrimp | 0.140 | 0.139 | (0.111,0.171) |
| | *Leptocottus armatus* | Pacific staghorn sculpin | 0.129 | 0.129 | (0.104,0.159) |
| | *Hemigrapsus oregonensis* | hairy shore crab | 0.126 | 0.125 | (0.101,0.155) |
| | *Ruditapes philippinarum* | Manila clam | 0.123 | 0.123 | (0.113,0.133) |
| | *Cymatogaster aggregata* | shiner perch | 0.123 | 0.122 | (0.098,0.151) |
| | *Cancer magister* | Dungeness crab | 0.121 | 0.120 | (0.096,0.149) |
| | *Batillaria attramentaria* | mud snail | 0.119 | 0.119 | (0.095,0.147) |
| | *Mya arenaria* | soft-shell clam | 0.119 | 0.119 | (0.096,0.146) |
| Slough | *Hemigrapsus oregonensis* | hairy shore crab | 0.425 | 0.424 | (0.312,0.543) |
| | *Crangon franciscorum* | sand shrimp | 0.324 | 0.322 | (0.241,0.412) |
| | *Mya arenaria* | soft-shell clam | 0.252 | 0.249 | (0.162,0.355) |
| All | *Cymatogaster aggregata* | shiner perch | 0.148 | 0.147 | (0.119,0.181) |
| | *Leptocottus armatus* | Pacific staghorn sculpin | 0.140 | 0.140 | (0.113,0.17) |
| | *Cancer magister* | Dungeness crab | 0.123 | 0.122 | (0.098,0.15) |
| | *Ruditapes philippinarum* | Manila clam | 0.122 | 0.122 | (0.112,0.132) |
| | *Mya arenaria* | soft-shell clam | 0.122 | 0.121 | (0.098,0.149) |
| | *Hemigrapsus oregonensis* | hairy shore crab | 0.117 | 0.116 | (0.094,0.143) |
| | *Crangon franciscorum* | sand shrimp | 0.115 | 0.115 | (0.092,0.141) |
| | *Batillaria attramentaria* | mud snail | 0.114 | 0.113 | (0.091,0.14) |

**S8 Table. The ranked relative abundance of each calibrated prey item by site type**, according to median sequencing read abundance (“Read abundance”) and median proportion of DNA in the stomach content sample (“True DNA abundance”; see Figure S4). ↑ Indicates an increase in rank, and ↓ indicates a decrease in rank.

| Rank | Read abundance | True DNA abundance |
|---|---|---|
| 1 | *C. franciscorum* | *C. franciscorum* |
| 2 | *L. armatus* | *L. armatus* |
| 3 | *C. aggregata* | *R. philippinarum* ↑ |
| 4 | *M. arenaria* | *C. aggregata* ↓ |
| 5 | *B. attramentaria* | *B. attramentaria* |
| 6 | *M. (Cancer) magister* | *M. (Cancer) magister* |
| 7 | *R. philippinarum* | *H. oregonensis* ↑ |
| 8 | *H. oregonensis* | *M. arenaria* ↓ |

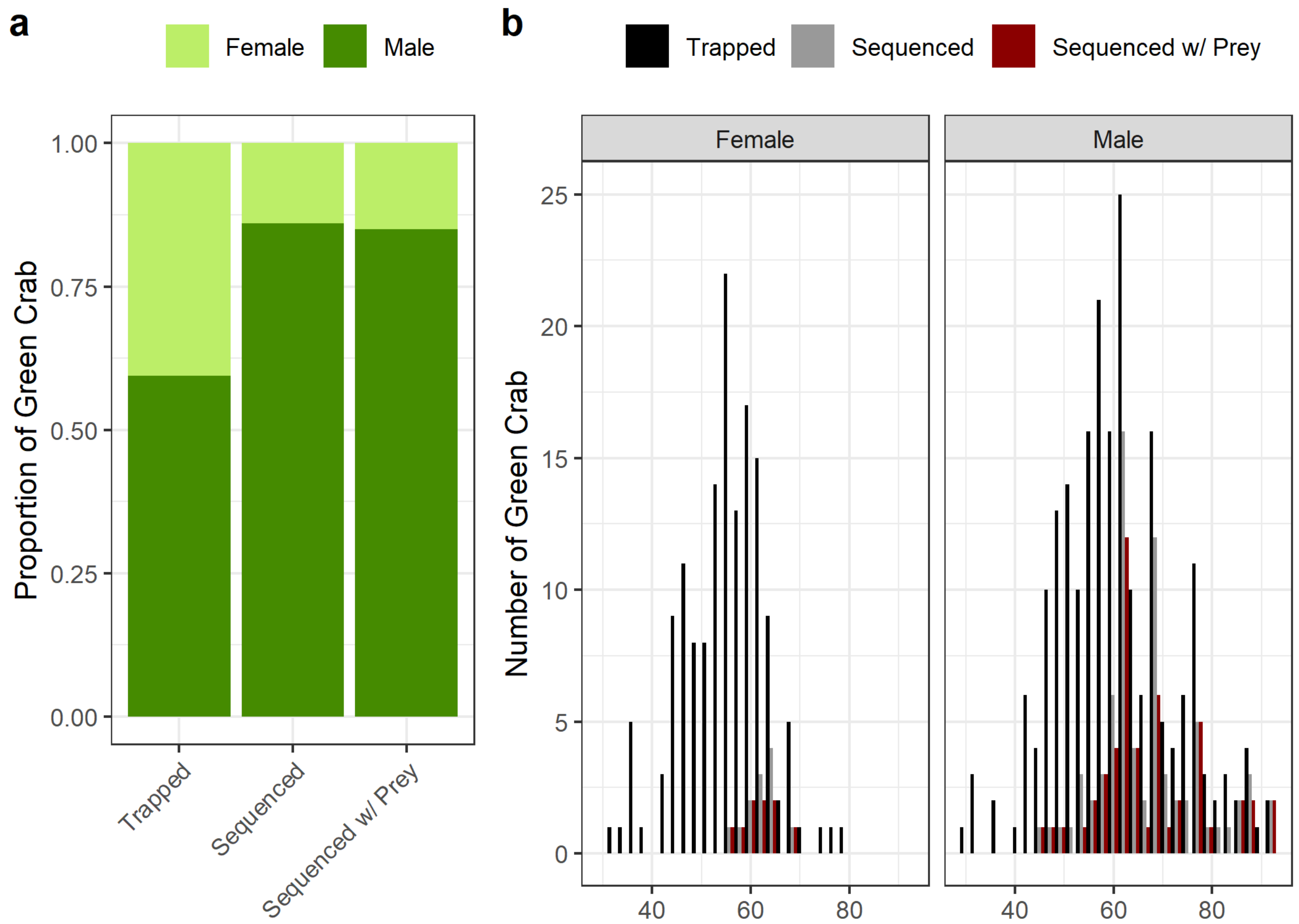


**S1 Fig. Sampling metadata according to sex**, including (a) proportion of crab at each step in Fig 1b belonging to each sex, and (b) distribution of carapace widths.

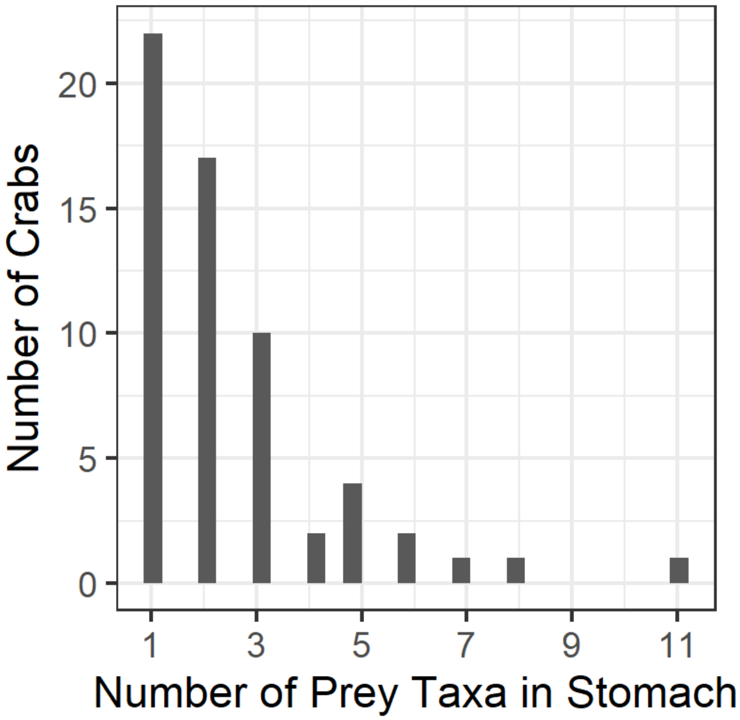


**S2 Fig. Distribution of α diversity of prey per crab stomach.**

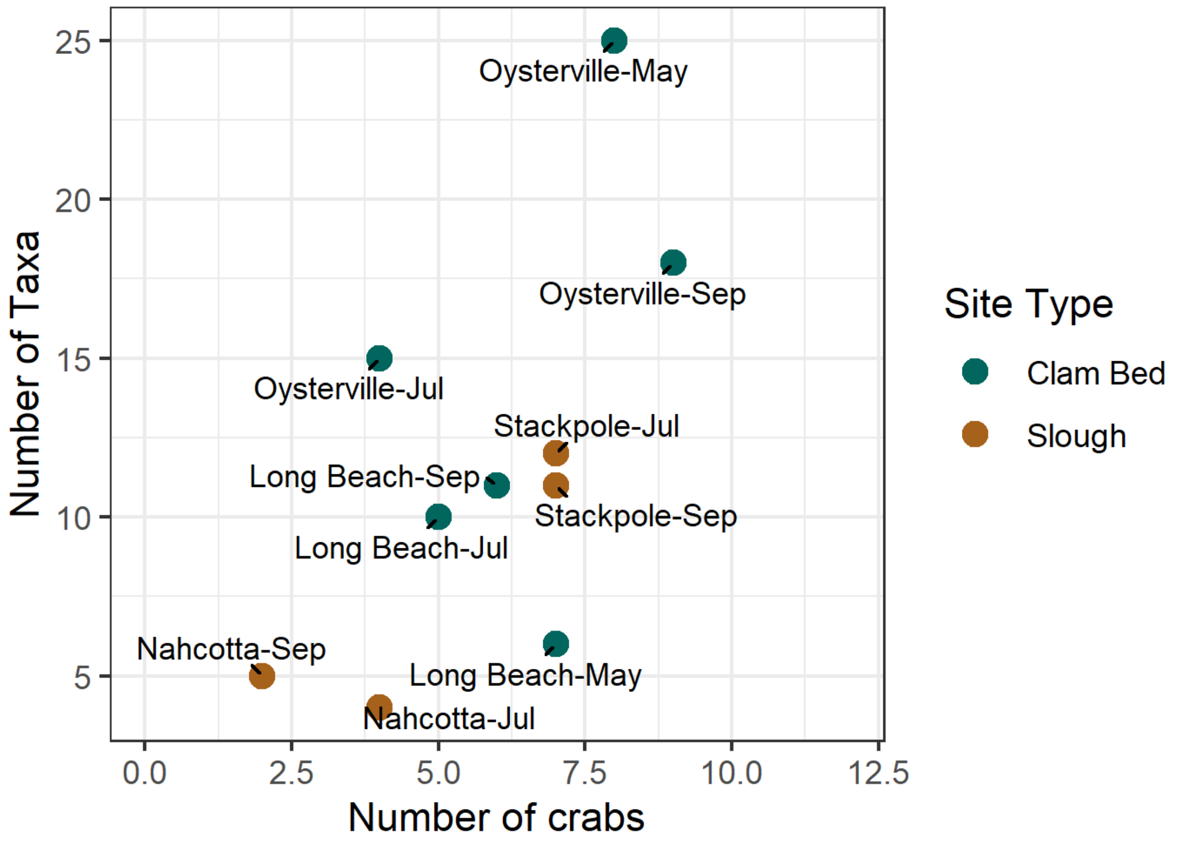


**S3 Fig. α diversity of prey per site / collection month, against the number of crab sampled.**

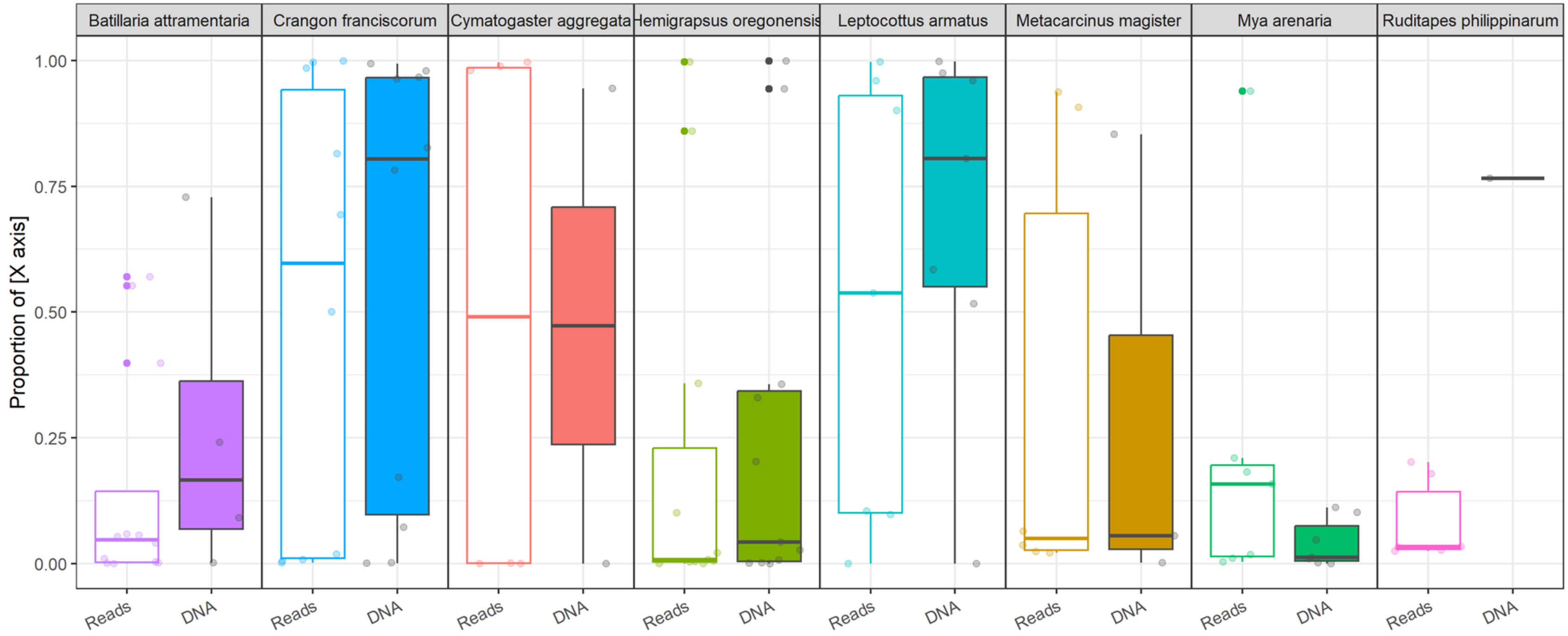


**S4 Fig. The proportion of reads ("Reads") and proportion of DNA ("DNA") representing each of the eight calibrated prey species in crab stomachs where the given species was detected.** Differences in the distribution of the proportion of reads and proportion of DNA for each species are a result of accounting for amplification efficiencies using the Shelton et al. (2023) quantitative model. Partially transparent points represent individual observations; for reads, these are laboratory samples (the technical replicates of the crabs included in the analysis), and for DNA, these are crabs; for example, note that the Manila clam was detected in one crab, but we sequenced six technical replicates of that individual. When crabs with a given calibrated prey species had different numbers of technical replicates, we randomly sampled read proportions to the least number of technical replicates for crabs in that group.

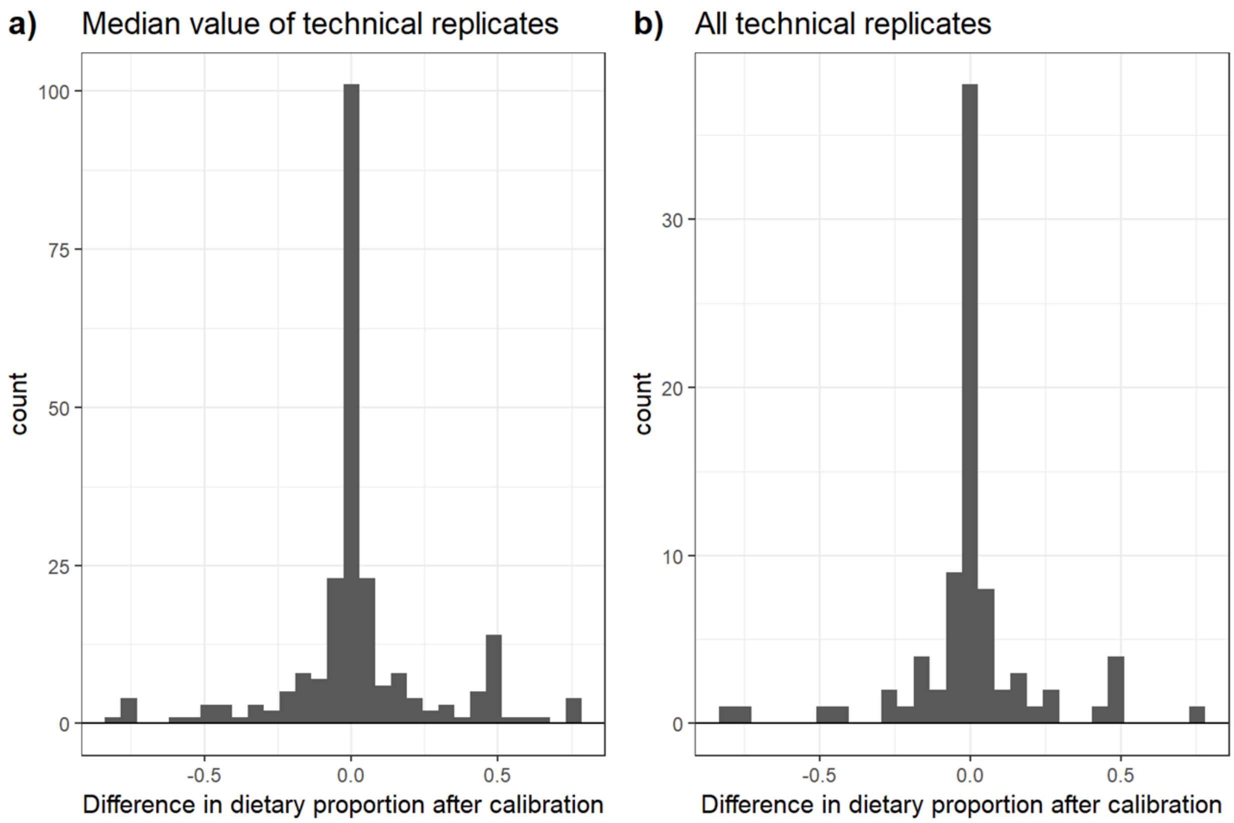


**S5 Fig. Differences in each species' dietary proportion after calibration**. For each of the eight calibrated prey species, the observed proportion of sequencing reads (uncalibrated) was subtracted from the estimated proportion of DNA (calibrated), by either (a) using the median proportion of sequencing reads across all technical replicates for an individual crab, or (b) subtracting the estimated proportion of DNA from the observed proportion of sequencing reads for each individual technical replicate.

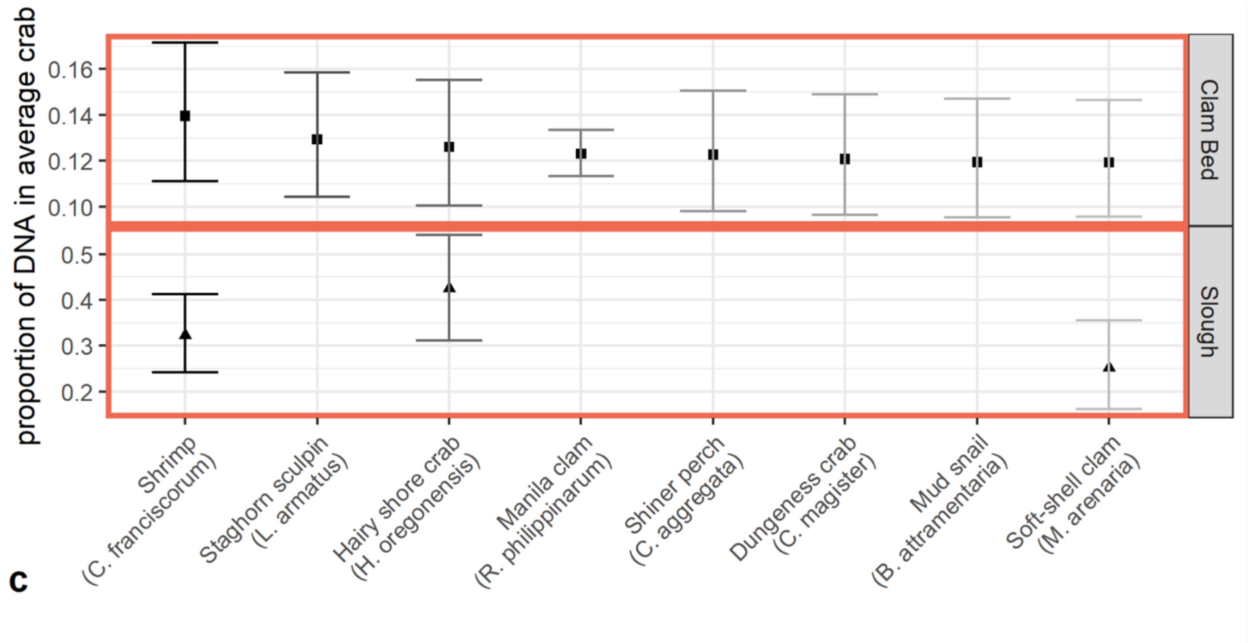


**S6 Fig. Mean estimated proportions of DNA contributed by calibrated prey species to an "average" crab diet, with confidence intervals.** "Average" diet shown separately for a green crab from clam bed sites and slough sites.
